# Muscle activation prediction in essential tremor through neuromusculoskeletal digital twinning and deep neural networks

**DOI:** 10.64898/2025.12.17.693373

**Authors:** Nuria Balbás, David Rodriguez, Filipe Oliveira-Barroso, Pablo Lanillos

## Abstract

Essential tremor (ET) is the most common movement disorder in adults, affecting up to 5% of the population over 65 years of age. Accurately predicting the dynamics of ET for each individual is crucial for optimizing therapies, such as sub-motor threshold stimulation (delivery of electrical currents below motoneuron activation), where the timing of stimulation is key for effective tremor reduction. Although there have been some efforts to implement machine learning predictive models, real-time prediction and estimation of muscle activation is still challenging due to the closed-loop nature of neuromuscular control, sensor noise, signal transmission delays, and scarcity of data. Here, we describe how a digital twin of ET—a computational neuromusculoskeletal model of ET deployed in SCONE simulator—allows for properly training deep recurrent neural networks (RNN) to predict muscle activation. Moreover, it permits parametrized synthetic simulation of the tremor. Results on predicting muscle activation from wrist flexo-extension movement show that the RNN has an average prediction accuracy of 81% and 83% with Long Short-Term Memory (LSTM) and Gated Recurrent Unit (GRU) gated neurons, respectively^3^. While this work still uses only synthetic data, it shows the potential for treatment optimization and personalized therapeutic strategies, such as peripheral electrical stimulation.

## 1 Introduction

Recent advances in digital twins, artificial intelligence and biomechanics software have opened up doors to new opportunities in the medical device industry, serving as tools to improve, test and design new treatments and therapies [1, 4]. On the one hand, digital twinning in movement disorders, besides allowing simulating and analyzing the pathology under various conditions, enables the generation of synthetic data that can be used to improve data-driven AI methods for optimizing and personalizing therapeutic strategies by using tools such as OpenSim [2]. On the other hand, recent therapies in peripheral electrical stimulation, such as sub-motor threshold stimulation in ET [5], require accurate temporal prediction of muscle activation to provide the correct stimulation and maximize tremor reduction. However, real-time prediction and estimation of muscle activation is still challenging due to the closed-loop nature of neuromuscular control, sensor noise, signal transmission delays, and scarcity of the data. While RNNs have been used for kinematic prediction [6], the present study focuses on predicting muscle activations.

This work describes how neuromusculoskeletal models can be used to generate heterogeneous synthetic data for training a deep artificial neural network to predict muscle activation. We modeled neuromusculoskeletal model of ET using a Central Pattern Generator (CPG) and spinal-muscle reflexes deployed into the SCONE simulator to generate tremor and to obtain data for training an RNN and allowing 1-second prediction of muscle activations given the wrist angle.

### 2 Materials and Methods

A muscle activation prediction pipeline (see Fig. 1) was developed, including a digital twin composed of a musculoskeletal model restricted to 1DoF (wrist flexo-extension) controlled by a neural oscillator (Matsuoka’s oscillator) in combination with muscular reflexes. The digital twin was used to generate synthetic data with frequency and amplitude variability to train a RNN with two different gating units (LSTM and GRU), with the aim of predicting one second of future muscular activations based on 600ms of kinematic wrist movement.

**Fig. 1.**
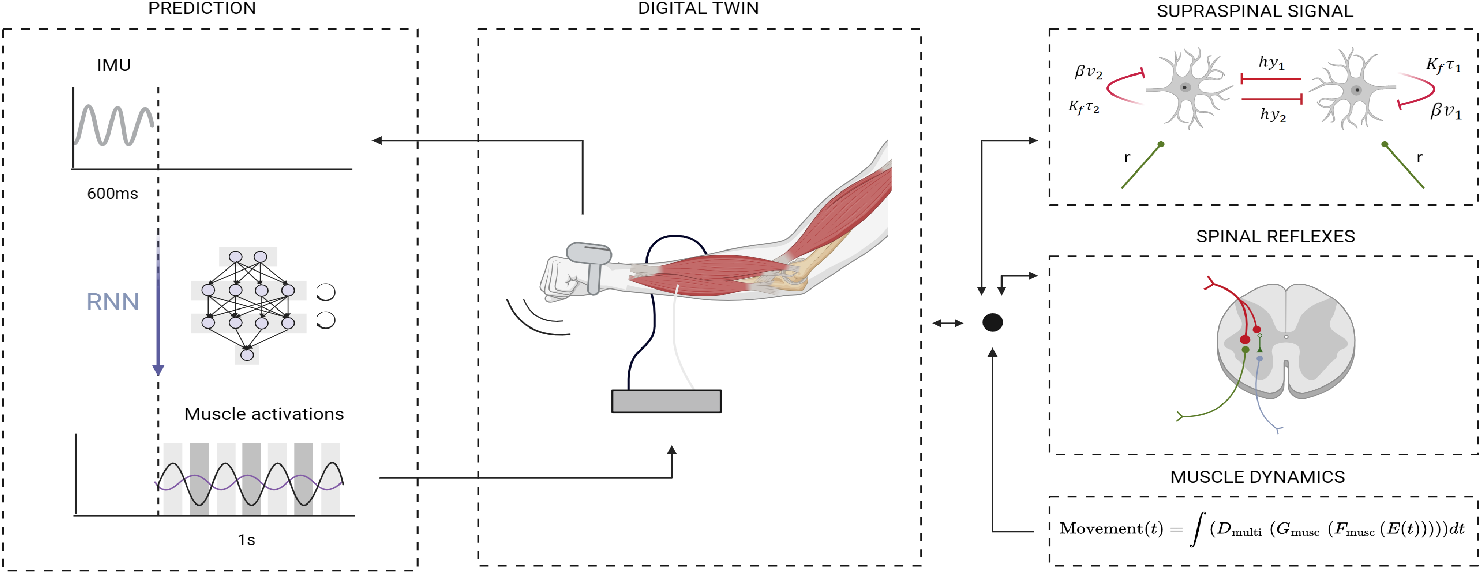
Proposed approach for muscle activation prediction in ET: RNN predictor and the digital twin with the musculoskeletal simulator and tremor computational model.

### 2.1 Digital twin

A simplified version of the MoBL-ARMS model [9] was implemented using Open-Sim and exported to SCONE for simulation. The musculoskeletal model was constrained to a single degree of freedom (DoF) at the wrist (flexion-extension), and was limited to five primary muscles involved in wrist motion. The forearm was positioned horizontally with the palm facing downward to emulate postural tremor conditions. To simulate neural rhythmic input, a CPG was implemented as a Matsuoka oscillator, based on recent computational tremor modeling [7]. To incorporate the influence of muscle force, muscle fiber length and muscle fiber velocity, we included spinal-muscle reflexes [3].

Within the sconepy package (a python wrapper of SCONE simulator), 7,000 independent simulations were generated. We introduced tremor variability through randomized base frequencies, oscillation amplitudes, and noise. Each simulation lasted 410 milliseconds, with a sampling rate of 100Hz, and employed distinct seeds and initial conditions to ensure heterogeneity and reproducibility. Outputs included both joint kinematics and muscle activations, stored in a database for offline training. From the collected simulation data, the wrist joint angle was used as model input, whilst the activations of the Extensor Carpi Radialis (ECRL) and Flexor Carpi Radialis (FCR) muscles served as prediction targets. Data was normalized by using a MinMaxScaler, fitted exclusively on the training subset to avoid data leakage. The data was then split by simulation (avoiding overlap) into 70% training, 15% validation and 15% test.

### 2.2 RNN predictor

Two types of sequence-to-sequence models were explored and implemented using PyTorch: a RNN based on LSTMs, and an RNN based on GRUs. The hyperparameters for both architectures were optimized using Optuna, for the learning rate (0.001 to 0.05), optimizer type (Adam, RMSEprop and SGD), number of layers (1 to 3), number of hidden states (20 to 50) and number of batches (16 to 64) in 50 trials with 10 epochs each. RNN predictors were trained using mean squared error (MSE) as the loss function, and the Adam optimizer with a learning rate of 0.001. Training was set to 70 epochs, with early stopping (patience= 10) to avoid overfitting. Training was accelerated via GPU when available.

## 3 Results

The RNN predictor performance was assessed using mean absolute error (MAE), root mean squared error (RMSE), and coefficient of determination on all dataset partitions. Additional qualitative assessments were also done to obtain graphs of the predictions, absolute errors per muscle in the form of histograms and mean prediction error per timestep. Figure 2 shows the input and prediction results for a single tremor sample from the test set. A data segment containing 600 ms of wrist flexion-extension angles is fed into the deep RNN, which outputs the normalized muscle activations for one second.The simulation data contained different base frequencies for each of the simulations. Three samples of different Fast Fourier Transforms (FFT) from the wrist velocity and position for three of the simulations can be found in Figure 3A. Furthermore, the intermuscular phase difference between the two antagonist muscles predicted (ECRL and FCR), was observed to have a mean and standard deviation within the range of experimental intermuscular phase difference of ET patients, as shown in Figure 3. Table 1 shows the RNN performance by comparing the best performing LSTM with the best performing GRU model in the prediction of muscle activation, obtaining better performance of the GRU model according to the MAE by 0.0007, the RMS Error by 0.0015, and the R^2^ by 0.016.

**Table 1.** Metrics results for the LSTM and GRU models.

| | Dataset | MAE | RMSE | $R^2$ |
| --- | --- | --- | --- | --- |
| LSTM | Train | 0.0065 | 0.0100 | 0.8246 |
|  | Validation | 0.0066 | 0.0102 | 0.8199 |
|  | Test | 0.0065 | 0.0101 | 0.8196 |
| GRU | Train | 0.0057 | 0.0085 | 0.8402 |
|  | Validation | 0.0057 | 0.0085 | 0.8372 |
|  | Test | **0.0058** | **0.0086** | 0.8356 |

**Fig. 2.**
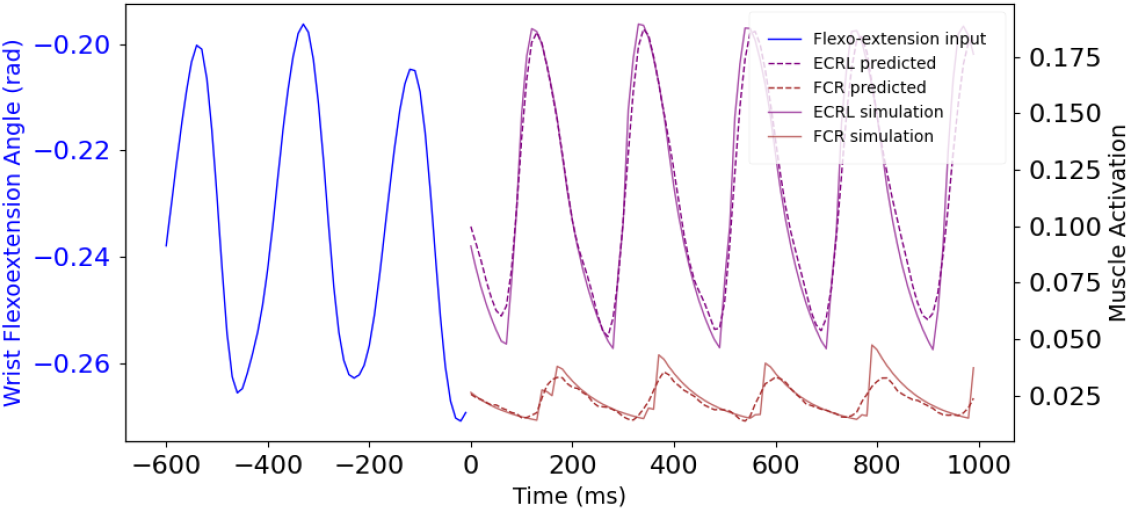
Kinematic input (blue line) and two muscle activation (ECRL and FCR) predictions from the LSTM network vs ground-truth muscle activations from the simulation. Extension is represented with an increase towards positive values, whereas flexion is represented with a decrease towards negative values.

**Fig. 3.**
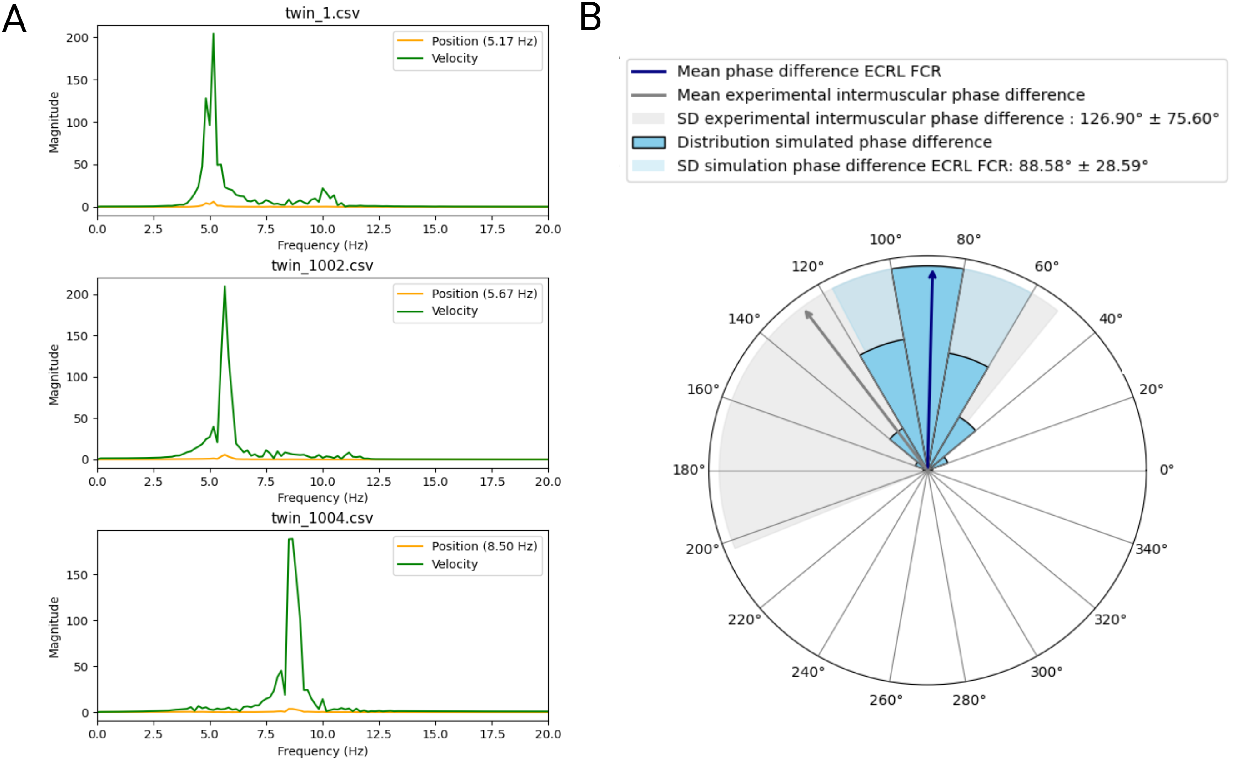
Analysis of generated tremors. (A) FFT of 3 different simulations from (wrist velocity and position). (B) Circular histogram of the instantaneous phase difference between FCR and ECRL muscle activation compared with experimental antagonist intermuscular phase differences in ET patients obtained from [8].

The value obtained by R^2^ further indicates that the 83.56% the predicted muscle activation dynamics can be explained by the GRU, and 81.96% by the LSTM.

The learning curves for both the LSTM and GRU RNNs from Figure 4 show no indices of overfitting, with higher learning rates in the initial epochs, and a stabilization in the epochs leading to early stopping. We further analyzed the statistical response for the whole prediction window. Fig. 4 bottom shows the mean error and standard deviation for the test samples. The ECRL muscle activation error shows a sustained average error of 0.02 but an increase in the standard deviation with time. This error accumulation indicates that the predictor behaves better closer to the initial timesteps.

**Fig. 4.**
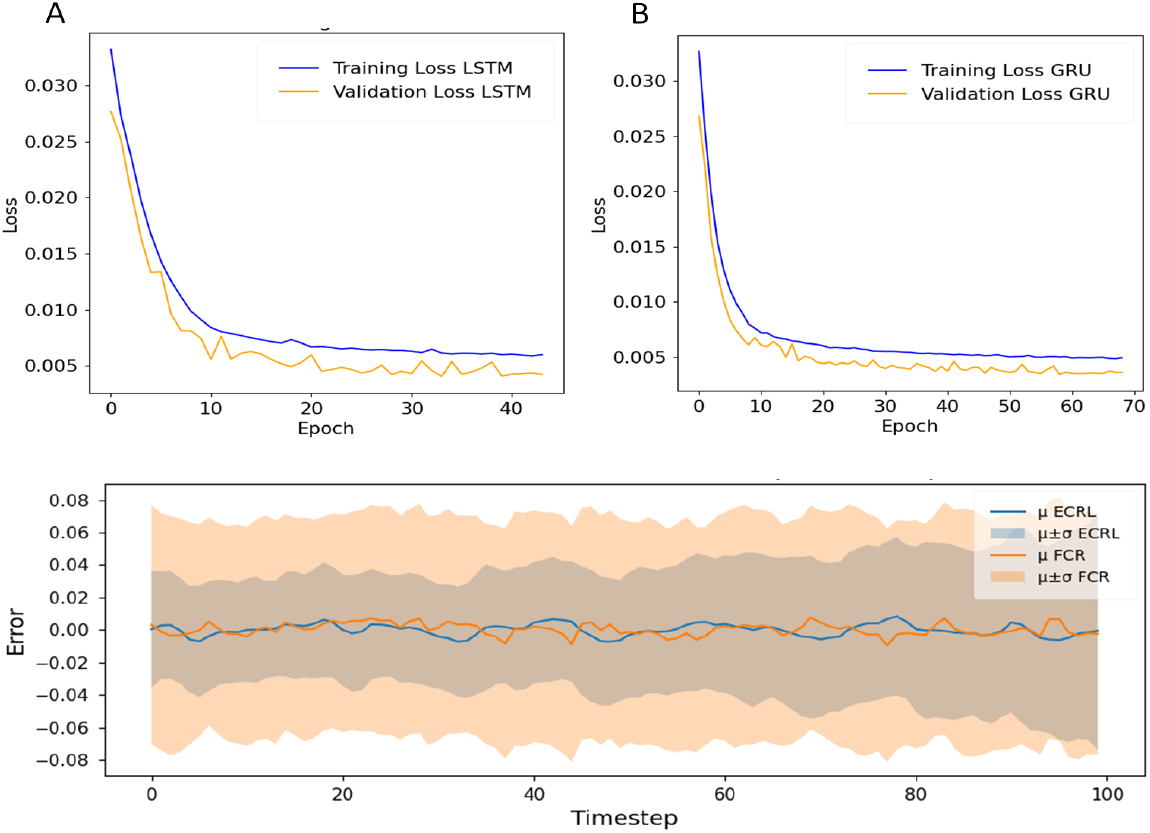
Training and average prediction accuracy per time step. (Top) Learning curves for best-performing (A) LSTM and (B) GRU RNNs, based on the MSE loss from normalized muscle activation. (Bottom) Muscle activation mean error and standard deviation per timestep for the LSTM network.

## 4 Conclusion

This study presents a neuromusculoskeletal digital twin framework for muscle activation prediction in ET by integrating biomechanical simulations and data-driven machine learning. A comprehensive simulation dataset was generated and used to train RNNs. Both LSTM and GRU-based models showed the ability to accurately predict muscle activity from wrist kinematics, achieving low inference times for real-time deployment. Ultimately, this work contributes to developing flexible frameworks for tremor modeling and prediction, which can be extended to other neurological tremor types and therapies, such as peripheral stimulation (e.g., sub-motor threshold stimulation) and robotic assistance. The modularity of the presented digital twin allows for future integration and validation with real-world IMU/EMG databases, extension to multi-DOFs arm models, the study of the influence of different positions and movements, and personalized calibration for clinical applications, as well as better control over the conditions in which the predictors are more accurate.

## Acknowledgements

This research was funded by the Spanish State Research Agency Ramon y Cajal Grant (RYC2021-031561-I) and Momentum Program (MMT24-CINC-01).

## Footnotes

3 The code and the data can be found in github.com/nuribal123/TremorNAIR.git.

